# Fossils improve phylogenetic analyses of morphological characters

**DOI:** 10.1101/2020.12.03.410068

**Authors:** Nicolás Mongiardino Koch, Russell J. Garwood, Luke A. Parry

## Abstract

Fossils provide our only direct window into evolutionary events in the distant past. Incorporating them into phylogenetic hypotheses of living clades can help elucidate macroevolutionary patterns and processes, such as ancestral states and diversification dynamics. However, the effect fossils have on phylogenetic inference from morphological data remains controversial. Previous studies have highlighted their strong impact on topologies inferred from empirical data, but have not demonstrated that they improve accuracy. The consequences of explicitly incorporating the stratigraphic ages of fossils using tip-dated inference are also unclear. Here we employ a simulation approach to explore how fossil sampling and missing data affect tree reconstruction across a range of inference methods. Our results show that fossil taxa improve phylogenetic analysis of morphological datasets, even when highly fragmentary. Irrespective of inference method, fossils improve the accuracy of phylogenies and increase the number of resolved nodes. They also induce the collapse of ancient and highly uncertain relationships that tend to be incorrectly resolved when sampling only extant taxa. Furthermore, tip-dated analyses which simultaneously infer tree topology and divergence times outperform all other methods of inference, demonstrating that the stratigraphic ages of fossils contain vital phylogenetic information. Fossils help to extract true phylogenetic signals from morphology, an effect that is mediated by both their unique morphology and their temporal information, and their incorporation in total-evidence phylogenetics is necessary to faithfully reconstruct evolutionary history.

## Introduction

Phylogenies underpin our ability to make sense of life on Earth in the context of its shared evolutionary history. In the absence of phylogenetic hypotheses, we would be unable to explain the myriad of biological phenomena that arise as the result of common ancestry, such as shared morphological features among seemingly disparate taxa. Many analyses that seek to investigate evolutionary events that occurred in the distant past do so using data from only living organisms. However, this might not be enough to faithfully recover evolutionary events that occurred in the deep past, as extant-only trees often lack the amount of information necessary to distinguish between alternative scenarios^1^ and can even favor incorrect results^2^. The incorporation of fossils into comparative analyses has been shown to have a positive impact on the inference of modes of macroevolution, ancestral character states and patterns of speciation and extinction^1–4^. Placing fossils in a phylogenetic framework is therefore necessary not only to understand their affinities, but can also be crucial to obtain an accurate picture of evolutionary history.

The effect that adding fossils has on phylogenetic inference remains equivocal however. Although fossils were originally dismissed as being too fragmentary to modify inferred relationships among living species^5^, a number of empirical studies have accumulated that show a common pattern of increased congruence between morphological and molecular phylogenies when paleontological data are introduced^6–8^. A recent analysis of multiple empirical datasets showed that adding fossil taxa to morphological matrices reshapes phylogenies in a manner that is entirely distinct from increasing the sampling of extant taxa^9^, a result largely attributable to the possession of distinctive character combinations in fossil taxa. However, given that the true tree of life is unknown, this study was unable to determine if these topological changes resulted in phylogenies that are more accurate.

Distinct combinations of morphological characters are not the only potential source of phylogenetic information that fossils can provide. The stratigraphic sequence of taxa in the rock record has also been proposed as a source of data with which to infer phylogenies. Although this idea was originally formalised in a controversial parsimony framework known as stratocladistics^10^, it has become popular again through the development of Bayesian tip-dated methods^11^. In this new framework, tree topology and divergence times are simultaneously estimated using the ages of fossil terminals to calibrate a morphological and/or molecular clock. This approach has been expanded with mechanistic models that include parameters for the rates of speciation, extinction and fossil sampling in an attempt to model the macroevolutionary process that generated the data, i.e. the fossilized birth-death process (FBD)^12^. Although the estimation of evolutionary timescales was the motivation for the development of these methods, they have also been found to strongly modify inferred topologies^13^, thus reshaping our understanding of evolutionary relationships. The use of temporal information from the fossil record as data in phylogenetic inference has been criticised however, with concerns ranging from the incompleteness of the geological record^14^ to the non-clocklike evolution of morphology^15^.

Inferring the phylogenetic position of fossils can only be done using datasets of morphological characters. Methods for inferring phylogenies from morphology have been increasingly scrutinised in recent years, establishing Bayesian inference as a valid alternative to parsimony approaches^16–18^. However, much of the early research on this topic has been criticized for simulating data under similar models of morphological evolution used in Bayesian inference, thus potentially biasing results^19^. In a similar vein, another study recently found tip-dated trees of extinct clades to be superior to undated ones^20^, a conclusion drawn from topologies simulated under the same birth-death processes used for tip-dated inference. Empirical analyses suggest fossils may overcome many limitations of morphological data for inferring the tree of life, but none of the above simulation studies explored the topological effects of sampling extinct lineages - or of the missing data typically associated with these.

In this study we employ a simulation approach^17^ to obtain character datasets and associated phylogenies that does not rely on any model later employed in the process of phylogenetic reconstruction, which we argue is necessary for results to be unbiased and extrapolatable to empirical datasets. We fine tune our simulations to produce mixtures of living and extinct taxa, as well as trees and characters that are empirically realistic based on comparisons with published datasets. Furthermore, we also mimic the presence of an ancient and rapid radiation^21^, comparable to the early radiation of placental mammals^22^ or the origin of most modern phyla during the Cambrian explosion^23^, as it is often suggested that fossils might be especially beneficial for resolving these kinds of radiations^24^. With these datasets we explore for the first time the topological effects of sampling different proportions of fossil taxa across a range of conditions of missing data, and investigate the relative behavior of tip-dated inference under the FBD prior (henceforth ‘clock’) relative to traditional undated approaches, i.e. maximum parsimony (MP) and Bayesian inference (BI).

## Results

The results of each analysis were summarized using standard consensus approaches and levels of precision and accuracy were calculated using both bipartitions and quartets. We define topological precision as the number of resolved bipartitions/quartets, and topological accuracy as the proportion of these that are correct (i.e., present in the true tree). Including fossil terminals in phylogenetic analyses increases the accuracy of phylogenetic reconstruction across all inference methods (Fig. 1). With no missing data, quartet-based accuracy consistently improves with an increasing proportion of fossils. Measured through bipartitions, accuracy is generally highest when 50% of terminals are fossils, and decreases with either higher or lower proportions of extinct taxa. Missing data generally limits the increase in accuracy provided by fossils, although the magnitude of this impact depends on the choice of method.

**Figure 1:**
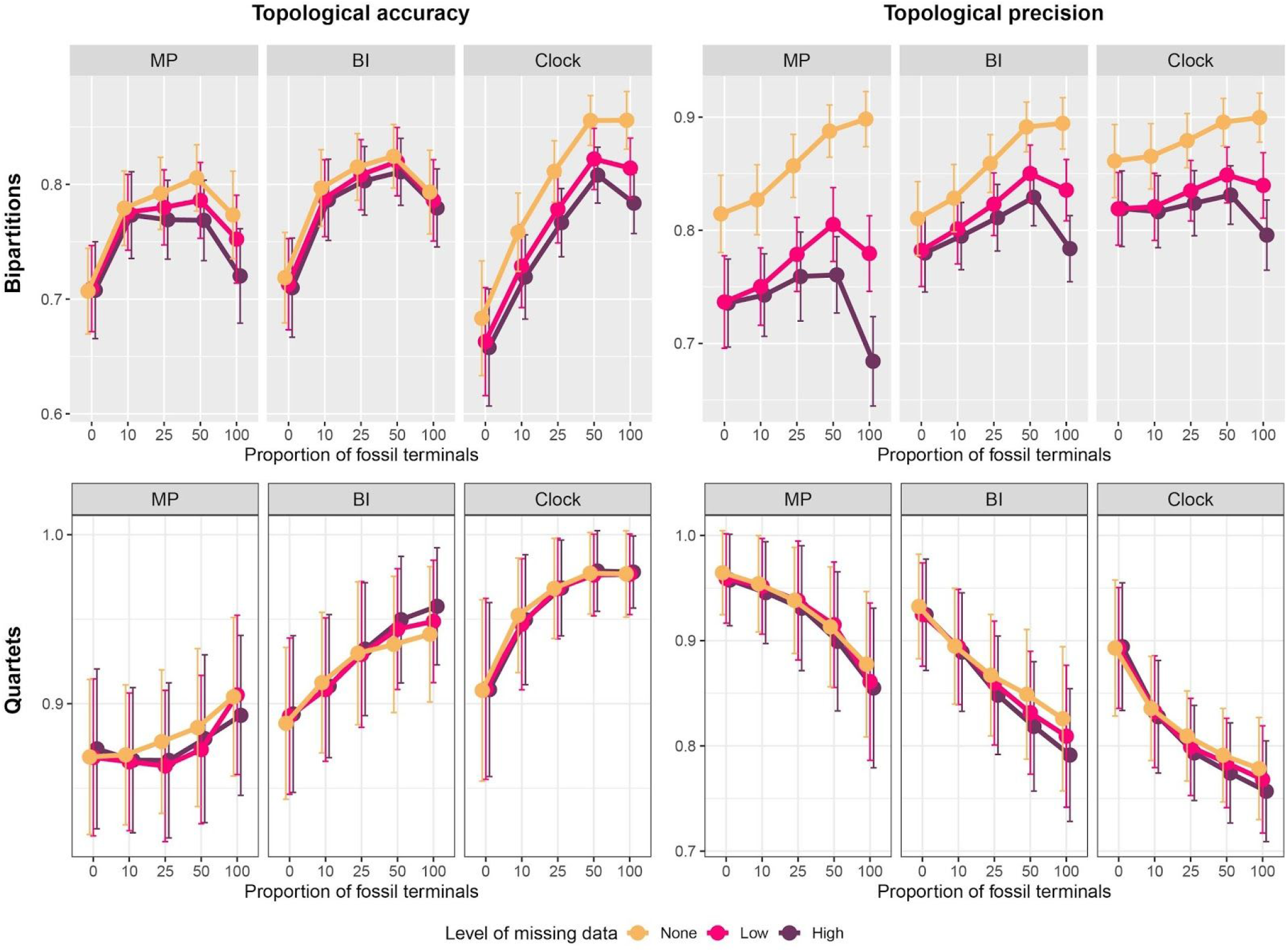
Impact of fossil sampling, missing data and method of inference on topological accuracy and precision. Accuracy (left) represents the proportion of resolved bipartitions/quartets, precision (right) the fraction of these that are correct. Values correspond to means +/− one standard deviation.

Topological precision increases with fossil sampling when measured using bipartitions, but decreases under quartets. This is because incorporating fossils tends to collapse a small number of deep nodes (which are present in a large proportion of quartets), while helping to resolve a larger number of shallower nodes (which collectively account for fewer quartets). This is supported by results shown in Supplementary Figure 1. Bipartition-based precision strongly depends on the amount of missing data, and MP is affected the most.

In order to explore how different methods of inference perform under the conditions explored, we summarized the overall differences between inferred and true trees using quartet distances (Fig. 2). This analysis reveals that the performance of both probabilistic approaches (both clock and BI) improves when living and fossil taxa are combined, relative to their behavior when datasets are composed entirely of either type of terminal. MP on the other hand remains unaffected by the proportion of fossils when no data are missing, but its performance declines with increasing proportions of incomplete fossils. Across all conditions explored, probabilistic approaches recover topologies more similar to the true tree than parsimony, a difference that widens with increasing missing data (Fig. S2). Analyses that enforce a morphological clock significantly outperform uncalibrated approaches when extinct terminals are incorporated (except where analyses are confined to fossil taxa with high levels of missing data). For any given level of missing data, the shortest distances between inferred and true topologies are always recovered when datasets combine fossil and extant terminals, and are tip-dated under the FBD model (Fig. 2). However, the relative benefit of tip-dating diminishes as the proportion of missing data increases (Fig. S2).

**Figure 2:**
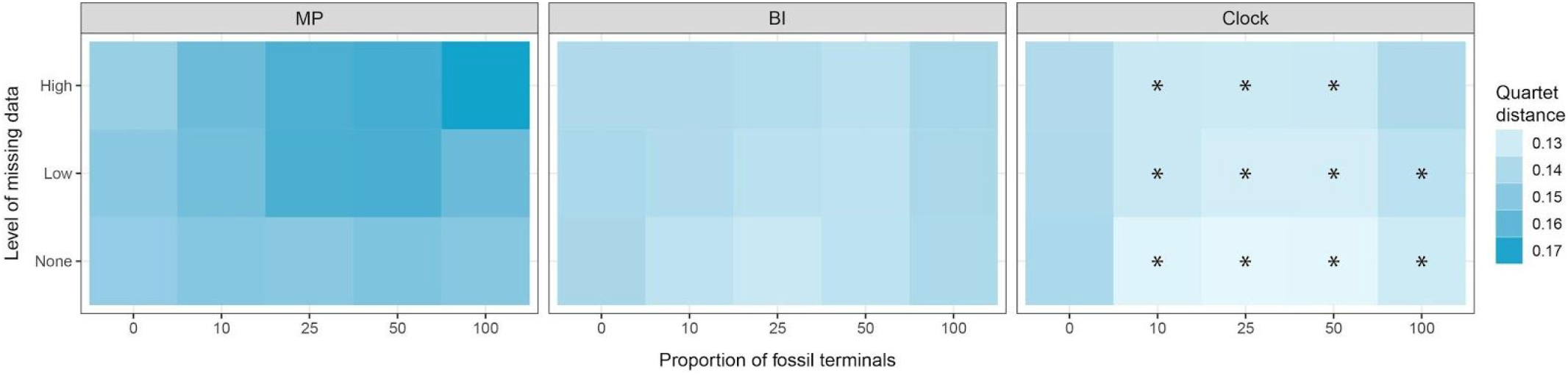
Average quartet distances between inferred and true trees across inference methods, proportions of fossil terminals and levels of missing data. For all conditions explored, clock analyses infer trees that are the most similar to the true trees (i.e., show the smallest distances), and MP the most dissimilar (i.e., show the largest distances). Asterisks mark the conditions under which the quartet distances of clock analyses are significantly lower than those of the second best method (i.e., BI).

To further characterize the topological changes induced by paleontological data, we explored whether the addition of fossils affects the ways in which relationships among extant terminals are resolved. Nodes connecting extant lineages were binned into three categories of equal duration (representing deep-, mid- and shallow divergences; see Fig. S3), and we recorded the proportion of nodes resolved correctly, incorrectly or left unresolved. Across inference methods, fossils help recover true relationships for mid and shallow divergences (Fig. 3). Fossils do not affect the proportion of correctly resolved deep nodes (including those involved in ancient and rapid radiations) and they increase the frequency with which these are left unresolved. This effect is especially strong in tip-dated inference, and seems to be a major reason why this approach performs best. However, fossil placement in tip-dated trees is less accurate than under undated BI, especially so for younger fossils (Fig. S4).

**Figure 3:**
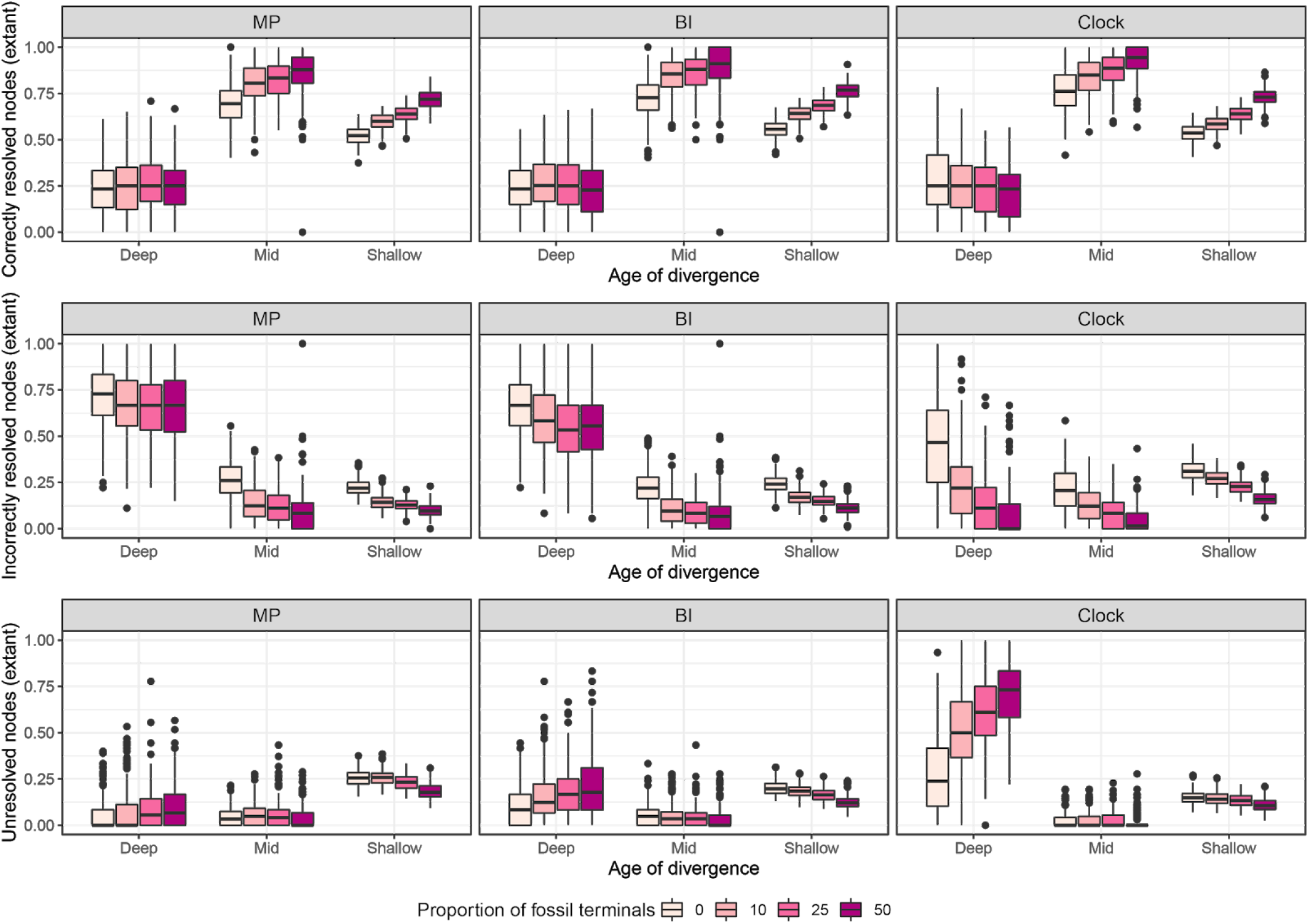
Effect of fossils on the way relationships among extant taxa are resolved (from top to bottom: correctly, incorrectly and unresolved). Results correspond to the high missing data condition, as this was designed to mimic empirically realistic datasets. Nodes connecting extant taxa were grouped into three time-bins of equal duration, representing deep-, mid- and shallow divergences.

## Discussion

Through simulations, we demonstrate that paleontological data (both morphological and stratigraphic) have a strong impact on tree inference across a wide range of realistic scenarios, supporting the results of empirical studies^6–9^. Analyses that incorporate paleontological data are more accurate than those based exclusively on extant taxa, regardless of inference method (Fig. 1). This improvement is partly driven by fossils helping to correctly elucidate relationships among living clades, especially among lineages separated by mid- to shallow divergences (Fig. 3). Hence we might expect the increased congruence between morphological and molecular trees found for some clades^6–8,25^ to reflect a general trend of consilience through improved accuracy as fossils are incorporated in phylogenetic reconstruction. Trees that combine living and extinct taxa also show a higher proportion of resolved nodes, while at the same time leaving more deep nodes unresolved (Figs. 1, 3). The phylogenetic analysis of morphological data has been previously shown to result in overprecise topologies^17,26^, a phenomenon we find to be mostly associated with deep divergences. With increasing fossil sampling, this overprecision is remedied as deep nodes collapse, increasing the accuracy of the resulting topologies. Therefore, fossils help recognize the high uncertainty associated with resolving such complicated phylogenetic questions with the use of small morphological datasets. Although missing data decreases both accuracy and precision, it has little effect on the resolution (or lack thereof) of deep nodes, and thus impacts quartet-based measures minimally (Fig. 1).

In recent years, probabilistic approaches that explicitly model the processes of morphological evolution and species diversification have increased in prominence. Our results corroborate recent studies^16–18^ that suggest probabilistic methods recover more accurate consensus topologies than MP (Fig. 2). This pattern holds true across all conditions explored, but becomes stronger with increasing levels of missing data, which adversely impact parsimony more than probabilistic methods (Fig. S2). Even though Bayesian approaches have been criticised for their handling of incompletely-coded morphological characters ^27^, we find missing data to have a comparatively small effect on Bayesian consensus trees. In contrast, realistic levels of missing data in MP analyses completely negates the benefit of a more thorough fossil sampling.

Probabilistic methods of inference can also directly employ the morphological and stratigraphic information from fossils to inform divergence times, allowing for greater flexibility in the integration of molecular and paleontological data^28^. Such synthetic, total-evidence, approaches have provided unique insights into the origin and evolution of numerous lineages^29–32^. However, the ways in which morphological and stratigraphic information interact to determine tree topology is an active research area that is arguably in its infancy^13,20,33^. For example, while some improvements in tree topology have been found when fossil ages are accounted for^20^, temporal data can also override morphological signals in potentially detrimental ways^13,34,35^. Here we show that Bayesian tip-dated methods that make use of stratigraphic information significantly outperform undated methods across most of the conditions we explored, indicating that stratigraphic ages provide important phylogenetic information^10,13,36,37^. However, the relative benefit of tip-dating seems to diminish in the presence of realistic levels of missing data (Fig. S2), to the point that tip-dated topologies of entirely extinct clades are not significantly better than undated ones (Fig. 2). Topological changes induced by tip-dating fossil clades (e.g.^38,39^) should therefore be considered cautiously. Furthermore, tip-dated inference under the FBD struggles to infer the position of fossil terminals (as also shown by ^33^), and is the least accurate method for placing young fossils (Fig. S4).

Fossils often overturn relationships among extant taxa, even when highly incomplete^9^. The way they do so, however, depends on the relative age of extant clades: fossils increase accuracy of mid- to shallow nodes, while decreasing overprecision among deep nodes, including those involved in ancient radiations (Fig. 3). Several authors have hypothesized such radiations are cases where a strong contribution from fossils might be expected, as they represent the only taxa that can directly sample the radiation event, and have character states that are less burdened by subsequent evolutionary change ^24^. Our results suggest instead that such ancient events of rapid diversification may be out of the reach of morphological signal. However, even though fossils do not help resolve ancient nodes in phylogenies, they nonetheless play an important role by inducing the collapse of the nodes involved in these deep divergences, mitigating the misleading signal provided by sampling only extant taxa. This effect is stronger when their temporal information is incorporated, and is a major driver of the increased accuracy of tip-dated phylogenies.

Reconstructing evolutionary processes that occurred in the distant past benefits from integrating molecular and paleontological data^3,40,41^, a goal that is facilitated by total-evidence dated inference^11,28–33,35^. Within this framework, inferring more accurate morphological phylogenies is crucial, as morphology has been shown to impact tree topology^42^ and divergence-time estimates^35,43^ even when combined with genome-scale datasets. Our analyses show that this goal can be achieved through increased fossil sampling, as both the morphological and stratigraphic information from fossils positively impact tree topology. However, tip-dating is also sensitive to the presence of missing data, and thus combining fossils with more complete datasets from living relatives (morphological and/or molecular;^33^), is likely to result in the most accurate inference of phylogenetic relationships.

## Materials and Methods

We simultaneously simulated 250 phylogenies and character matrices using TREvoSim v2.0.0^17^, newly released with this paper [whilst this manuscript is in review, the v1.0.0 can be found <u>here</u>; v2.0.0 source code {for Linux} or releases {for Windows/MacOS} can be provided through the editor on request]. Simulation settings were chosen to emulate realistic properties (rates of evolution, distribution of branch lengths, levels of treeness - i.e. the fraction of total tree length that is on internal branches;^44^). This was based on comparisons with 12 empirical datasets (see SI, Table S1 and Figs. S5-6). The simulation settings also incorporate a series of deep and short internodes: a tree shape comparable to that of clades that underwent an ancient and rapid radiation^21–23^. Through subsampling, we built a total of 11,250 morphological matrices from these simulations, varying levels of missing data and proportions of fossil terminals. Imputation of missing data was designed to produce either low (25% for extant taxa, 37.5% for fossils) or high levels (25% and 50%, respectively); the latter condition designed to mimic realistic levels of missing data as found across empirical datasets. Multiple iterations of each condition further allowed us to factor out topological effects due to character and taxon sampling decisions. All analyzed matrices comprised 100 terminals and 300 parsimony-informative characters, and were analyzed using maximum parsimony (MP), and both undated (BI) and tip-dated (clock) Bayesian approaches. In the latter case we used either the fossilized birth-death^12^ or birth-death tree prior, depending on whether fossil taxa were sampled or not, respectively. Parameters for tip-dated analyses, such as prior distributions for the tree height and tip ages, were informed by data mined from paleontological databases (see SI). Inference was performed using TNT 1.6^45^ and Mrbayes 3.2^46^. For more details see SI.

We used standard consensus methods (i.e., strict consensus for MP; majority-rule consensus for probabilistic methods) to summarize the analyses, and compared the inferred consensus topologies to true (simulated) trees using bipartition and quartet-based definitions of precision and accuracy. The overall performance of alternative methods of inference under different conditions was summarized using quartet distances^16^ between estimated and true trees. TREvoSIM is an individual-based simulation with its own species definition: it does not rely on either Markov models for the generation of character datasets, nor birth-death processes for the simulation of topologies, and should therefore be unbiased towards alternative methods of inference. Significant effects were explored by comparing the distributions of quartet distances between the best (clock) and second best (BI) methods across conditions, using *t*-tests corrected for multiple comparisons.

We further assessed the interaction between paleontological data and methods of phylogenetic reconstruction by exploring the effects of incorporating fossils on the inference of relationships among living clades. To do so, we pruned consensus trees down to the subset of living terminals and recorded the proportion of correctly resolved nodes at different temporal depths (deep, mid and shallow, defined as time-slices of equal duration; Fig. S3). Furthermore, we also explored the different ways in which inference methods resolved the position of fossil terminals (i.e., correctly/incorrectly/unresolved) depending on their relative ages. All analyses were performed using data and code included in the supplementary files. Further details on data simulation, tree inference, and statistical analyses is placed in SI file xxx.

## Acknowledgements

We thank Derek E. G. Briggs for helpful comments that greatly improved the clarity of this manuscript, Martin R. Smith for help with the interpretation of quartet distances, Joe Keating for discussion regarding analyses, and Serjoscha Evers for providing a preliminary empirical dataset for turtles. The HTCondor service was provided by the IT Services Research Infrastructure team at the University of Manchester, and runs the HTCondor software developed by the CHTC Team at UW-Madison, Wisconsin. This work was supported by the Natural Environment Research Council [grant number NE/T000813/1 awarded to RJG].

## Supplementary Information for

### Detailed methods

#### Data generation

Much of the research on the relative performance of maximum parsimony and Bayesian inference as methods of phylogenetic reconstruction from morphology employed stochastic models to generate the simulated data (e.g., refs^18,26,26,47^). Where these are also the models of morphological evolution employed by software to infer trees under probabilistic methods, this decision has been criticised for potentially generating a bias towards favoring the use of Bayesian methods^19,48^. Another recent study found tip-dated trees of extinct clades to be superior to undated ones^20^. Extrapolating this finding to empirical data is made challenging by the fact that fossils lacked missing data, and that data were simulated using the same birth-death processes used in the process of inference. Given the impact of the tree prior on the topologies recovered^49^ this decision could potentially bias analyses towards favoring tip-dated inference, and it is likely that the history of diversification in real clades deviates from constant-rate birth-death dynamics^50,51^. Here, we generate phylogenies and morphological matrices in a way that does not rely on any model later employed in the process of phylogenetic inference, in order to minimise bias and maximise the extent to which we can extrapolate from this study to the analysis of empirical datasets.

Character datasets and associated phylogenies were created simultaneously using the individual-based simulation software TREvoSim, which has been fully described by Keating et al.^17^. In this program, individuals are represented using binary strings, which ultimately form the character data in the simulation. These strings are—through comparison with an environment—used to calculate individuals’ fitness. Those individuals compete against others within a structure called the playing field, and the fitness of an individual dictates the probability it will duplicate. The simulation employs a lineage-based species concept, and as it runs a tree representing the evolutionary relationships of species is recorded. This phylogeny, and associated character data, is output from a simulation once the requested number of taxa has evolved.

The data for the present study were generated in TREvoSim to meet a number of benchmarks based on the analysis of twelve empirical datasets, as outlined below. In order to achieve these properties—and as part of ongoing development of the package—a number of features have been added to TREvoSim, and are found in version 2.0.0, the release accompanying this paper. The simulation now allows multiple playing fields in which populations of digital organisms live and compete, which can have independent or identical environments. This allows a finer control of the number of extant terminals at simulation termination and the treeness of the resulting topologies (as a result, simulations can comprise several clades evolving independently). When an organism is duplicated, it is returned to the playing field, overwriting another individual: in release 1.0.0 this was the least fit member of the playing field. In release 2.0.0 there is an option to overwrite a random taxon. This allows a wider variance in terminal branch lengths. In order to provide finer control over the fitness landscape within the simulation, user control of the fitness target (i.e. the count of ones in the genome to environment hamming distance which is considered the peak fitness; see^17,52^) is now provided. Setting this to zero provides fewer fitness peaks: a count peaks functionality has been added to provide a histogram of possible fitnesses for a given set of masks and simulation settings. Finally the software now provides multiple environments against which the fitness of all organisms is assessed. Within a playing field, the fitness of any given organism is defined by the environment they are best suited to. This encourages numerous competing lineages in any given playing field, and creates more symmetrical trees. Additionally, the code has been refactored, the simulation now makes use of the Qt QRandomGenerator tools rather than incorporating random data, and a user-accessible test suite has been added. More details are provided in the release notes for the software.

For the current study, we generated data using simulations of 500 characters, that ran until the trees comprised 999 terminals (the maximum terminal count in TREvoSim). We employed 5 playing fields of size 40, each with 5 non-identical environments of 3 masks, a species difference of 8, an unresolvable cutoff of 2 (no identical terminals) and random overwrite enabled. All other settings employed defaults: we provide an SI file which will load these settings in TREvoSim (SI File XXX).

#### Data preparation

All subsequent steps of manipulation of topologies and character matrices were performed within the R environment^53^. TREvoSim runs that have multiple playing fields can produce trees with zero length branches, where speciations occur from the starting species in multiple playing fields within the same iteration. While this is not biologically implausible, it does present significant challenges for phylogenetic inference^21,54,55^, and results in reference topologies that are not fully bifurcating. Therefore, we ran many TREvoSim simulations and retained 250 simulations with no zero length branches for further study (SI file XX.r). These phylogenies contained a mean of 150 extant terminals at termination and exhibited a range of tree symmetries (Fig. S7), as estimated using Colles’ index^56^. They also incorporated an early radiation with short internodes as the lineages became established within each playing field, a tree shape broadly comparable to that of clades that underwent ancient and rapid radiations^21^, such as occurred during the Cambrian explosion^23^ and at the origin of the major lineages of placental mammals^22^ and insects^57^. Analysis of morphological coherence^58^ shows a linear correlation between morphological and patristic distances, with saturation occurring only between the deepest divergences (Fig. S8).

Prior to further analysis, we removed fossils at random until only 300 terminals remained, a step that emulated the reality that a significant amount of extinct biodiversity is not captured in the fossil record. This left datasets with an average of 150 extant and 150 extinct terminals, although values varied between a minimum of 136 and a maximum of 169 extant terminals (Fig. S7). Simulations were then compared to twelve empirical datasets (including morphological datasets and associated time-calibrated topologies) to corroborate the realism of their properties, following best practices for simulation studies in paleobiology^59^. Empirical datasets constitute a mix of tip-dated morphological or total-evidence trees, which either sample both living and extinct terminals, or only fossils (see Table S1). We compared: levels of morphological variability (measured as the number of parsimony steps per character); distributions of branch lengths; and values of treeness. The number of parsimony steps were tallied only for characters that were both binary and variable, as this is the type of data produced by our simulations, and were estimated using functions from R package *phangorn* ^60^. In all cases, we confirmed that the values exhibited by our simulations were comparable to the range of values displayed by empirical datasets (Figs. S5-6), and can thus provide insights on the behaviors of phylogenetic inference from morphological data.

Datasets were subsequently subsampled further to a size of 100 taxa and 300 characters, a final size that is shared by all of our analyses. Taxon subsampling was performed so that the final matrices contained different amounts of extinct lineages, including 0, 10, 25, 50 and 100% fossil terminals. Furthermore, the datasets were also subject to one of three different treatments of missing data, including no missing data, low levels of missing data (25% for extant taxa, 37.5% for fossils), or high levels of missing data (25% for extant taxa, 50% for fossils). The high missing data treatment is similar to the average values found across empirical datasets (proportions of missing data 28.9 and 54.2%, in extant and fossil taxa respectively; see Table S1), and can thus be considered the most realistic. As defined here, low levels of missing data represent an optimistic condition, observed under circumstances of exceptional preservation or in clades with unusually high levels of preservation. The chosen amount of missing data in fossils under this condition is, for example, similar to the amount of missing data present in the trilobite datasets (i.e., 36.3%). The condition of no missing data is explored to better understand the behavior of methods of phylogenetic inference under ideal conditions. Even though missing data is highly structured in empirical morphological datasets^61–64^, our simulated characters are independent and thus an approach to missing data imputation that implements some degree of correlation was not deemed necessary. Data was therefore deleted at random from the entire dataset, which also ensured some stochastic variability in the final proportions of data found among terminals of the same dataset. After taxon subsampling and missing data imputation, 300 parsimony-informative characters were selected at random.

In order to emulate the fact that different phylogenetic analyses of the same clade often sample different taxa and characters, this entire process was iterated three times for each combination of dataset, level of missing data and proportion of fossil terminals. This resulted in alternative datasets derived from the same evolutionary history, but differing in the traits and terminals that were incorporated. Overall, this procedure generated 11,250 final character matrices (250 simulations * 5 proportions of fossils * 3 levels of missing data * 3 iterations). Matrix and tree manipulations relied on custom R scripts (File SXXX) and made use of functions in packages *ape*^65^, *Claddis*^66^ and *phytools*^67^.

Even though we replicated many of the key features of empirical morphological datasets that we believe need to be considered in order to draw general conclusions regarding the behavior of methods of phylogenetic inference, we recognise - and would like to highlight - other aspects of our simulations that are likely not representative of those of empirical data. For example, even when missing data was applied at random to entire blocks of extant and fossil terminals, thus generating some level of variation in the total amount of coded data per taxon, missing data is likely to be much less homogeneously distributed across both taxa and characters in empirical datasets^61,62,64^. Furthermore, our simulated datasets do not take into account character contingencies, such as those generated by the hierarchical relationships that arise from the ontological dependences between morphological traits^68^, for example. Finally, most phylogenetic analyses sample taxa so as to capture all or most of the main lineages of the clade under analysis, an approach known as diversified sampling^69^. In our subsampling step that reduced datasets from 300 to the final 100 terminals included in phylogenetic analyses, and which simulates the step of taxonomic sampling made during matrix construction, we selected terminals at random.

#### Phylogenetic inference

All character matrices were analyzed using three methods of phylogenetic inference: maximum parsimony (MP), uncalibrated Bayesian inference (BI), and time-calibrated Bayesian inference under a morphological clock (Clock). Inference under MP was performed using TNT v. 1.5^45^ under equal weights, using driven tree searches with five initial replicates that were subject to new technology search heuristics^70,71^. Search was continued until minimum length was found twenty times. TBR branch swapping was then performed on the topologies in memory, retaining up to 50,000 maximum parsimony trees (a TNT batch script can be found as SI File XXX). Inference under BI was performed in MrBayes 3.2^46^ under the Mk + Γ model^72^ with correction for only including parsimony informative characters. Two runs of four Metropolis coupled MCMC chains were continued for either twenty million generations or until an average standard deviation of split frequencies (ASDSF) < 0.005 was attained. Previous analyses had continued analyses until ASDSF values of 0.01 were met, which was taken as indication of a thorough sampling of the posterior distribution of topologies^9,18^. The ASDSF values attained by our analyses were registered upon completion, revealing median values < 0.01 across all conditions for both tip-dated and undated Bayesian methods (Fig. S9). Given that we focused exclusively on topologies, we did not ensure other parameters converged. It should be noted that several parameters intrinsic to inference under the fossilized birth-death (FBD) process, including those involved in determining the clock rate and branch length distribution have not been found to affect overall topological accuracy of tip-dated analyses^20^. Trees were sampled every 1,000 generations and the initial 50% was discarded as burnin.

Clock analyses were also run in MrBayes, primarily under the FBD process but using the birth-death branch length prior when fossils were not sampled. In order to assign age uncertainty to fossils, the entire Paleobiology Database (PBDB; https://paleobiodb.org/) was downloaded and used to build a distribution of species’ longevities. Occurrences assigned to the same species were grouped, and longevities were defined as the time spanned between the minimum age of their first occurrence and the maximum age of their last one. The distribution of longevities was found to approximate an exponential distribution with a rate parameter of 0.115 (estimated using ‘fitdistr’ in *MASS*^73^), corresponding to a mean duration of 8.67 Ma. An exponential distribution also ensured short intervals were more likely to be drawn, while also allowing long durations to occur occasionally^74–76^. Time intervals were generated by sampling from this distribution and assigning to fossil terminals in such a way that the true tip age is contained randomly within this interval^20^. A minimum interval of 0.0117 Ma (the shortest one included in the PBDB) was enforced, and ranges were checked not to cross the present. Time ranges were treated as uniform priors for the age of fossil terminals, an approach that has been found to help recover correct topologies^20^. Given that the chronograms simulated by TREvoSim are scaled to the number of iterations, we translated this to absolute time by assuming a 210.9 Ma old root, given the average timespan of the 12 empirical datasets used as reference (see above), and rescaled branch lengths accordingly. This true root age was treated as an unknown parameter. To establish a tree age prior, we used an offset exponential distribution with a minimum value set to the minimum of the age range of the oldest preserved fossil (i.e., the oldest fossil in the full 300-taxon dataset; note that this taxon might or might not be a terminal in the analysis dependending on the random subsampling step). To establish a mean for the distribution, we scraped the fossil calibration database^77^ using functions from the *rvest* package^78^ and obtained 202 node calibrations with both minimum and maximum values. The average difference between these (107.7 Ma) was used to establish a soft maximum, setting the mean of the offset exponential distribution so that 95% of prior probability lies between the minimum and this value plus 107.7 Ma. The probability of sampling extant taxa was fixed to the true value, and the species sampling strategy was set to either ‘fossiltip’ or ‘random’, depending on whether analyses incorporated or not extinct terminals, respectively. The first of these ensured fossil taxa could not be recovered as sampled ancestors. Even though allowing for direct ancestor-descendant relationships is a major aspect of the realism of the FBD process^12,79^, TREvoSim simulations record the morphology of fossils only upon the extinction of the lineage. An independent gamma rate (IGR) prior was used for the morphological clock, the speciation probability prior was set to an exponential distribution with a rate of 10.0 and default values were used for all other priors and parameters. Two runs of four chains were continued for either fifty million generations or until reaching an ASDSF < 0.005. All other settings were as for the BI analyses. As shown in Fig. S9, these settings were enough to ensure topological convergence was attained.

### Statistical analyses

As done by previous studies (e.g.^19^), phylogenetic analyses were summarized using standard consensus approaches (i.e., strict for MP, 50% majority-rule for BI and clock), and the consensus topologies were compared to the corresponding true trees using a suite of approaches. Although differences in the behavior of inference methods have been found depending on whether optimal or consensus topologies are employed^17^, the amount of data generated here precluded the use of samples of optimal topologies. Furthermore, even when sets of optimal trees are (or should be) routinely employed to draw macroevolutionary inferences^80^, systematic studies generally rely only on consensus trees to draw their conclusions.

First, we measured the topological precision and accuracy of the analyses using both bipartitions and quartets. Topological precision was measured as the overall number of resolved bipartitions/quartets, topological accuracy as the proportion of these that are correct (i.e., present in the true tree). Furthermore, we used quartet distances as a summary statistic of the overall difference between true and inferred trees. Quartet distances have been found to outperform measures of tree similarity based on bipartitions^16^, especially when the topologies being compared are not fully bifurcating. Quartet distances are also less susceptible to wildcard taxa and tree shape, and have been found to be more precise and less prone to saturation, than symmetric distances based on bipartitions^16,81^. For these analyses, we first averaged the values obtained from different iterations of the same dataset under the same conditions, which just differed on taxon and character sampling. This gave us estimates of precision, accuracy and quartet distances that average out the effects of decisions taken during matrix construction, and should better approximate the overall difficulty of estimating relationships across the range of conditions explored. For quartet distances, the distribution of the best and second best methods of inference (clock and BI, respectively) were compared using *t*-tests, and *P*-values were corrected for multiple comparisons using the Benjamini & Hochberg^82^ correction.

A major theme in the discussion of the relevance of paleontological data for phylogenetic inference is whether fossils are able to improve our estimates of the relationships among living clades^5,9,24,83–85^. In order to explore this, we pruned down all inferred trees to just the extant taxa, and analyzed the impact that increasing fossil sampling has on the accuracy with which their relationships are reconstructed. In order to further account for a possible relationship with divergence times, we subdivided trees into three equal time bins representing shallow, mid and deep divergences (Fig. S3). Nodes connecting extant taxa in the true trees were then classified as being resolved correctly or incorrectly in the inferred consensus tree, depending on whether the node corresponding to the last common ancestor of taxa in that clade gave rise to a single clade of identical composition (regardless of its internal topology) or not. In case such a node constituted a polytomy (i.e., gave rise to more than 2 descendant lineages), all possible resolutions of this polytomy were explored, and the node was considered unresolved if a clade with the exact same composition as that of the true tree was found among the possible resolutions. If a clade with the correct composition could not be generated, the node was also counted as incorrectly resolved. Values were then expressed as fractions of the total number of nodes present within each time bin in the true tree. Analyses composed of only extinct terminals were excluded. Furthermore, we also assessed the way in which fossils of different ages are resolved across inference methods. Fossil terminals were classified as either being unresolved (i.e., attaching to a polytomous node), correctly or incorrectly resolved. We counted correctly resolved fossils as those whose sister group has the same composition in reference and inferred trees, regardless of internal relationships. We also subdivided the time spanned by reference trees into 20 time bins of equal duration, and estimated the proportion of fossils in each category for every bin. For both of these analyses (fossil placement and effect on extant nodes), we restricted the data to replicates with high levels of missing data, which is the condition most similar to empirical data (see above). Quantification of these metrics relied on custom R scripts (File SXXX) and made use of functions in packages *ape*^65^, *phangorn*^60^, *phytools*^67^, *Quartet*^86,87^ and *TNTR*^88^.

## Supplementary Tables

**Table S1:**
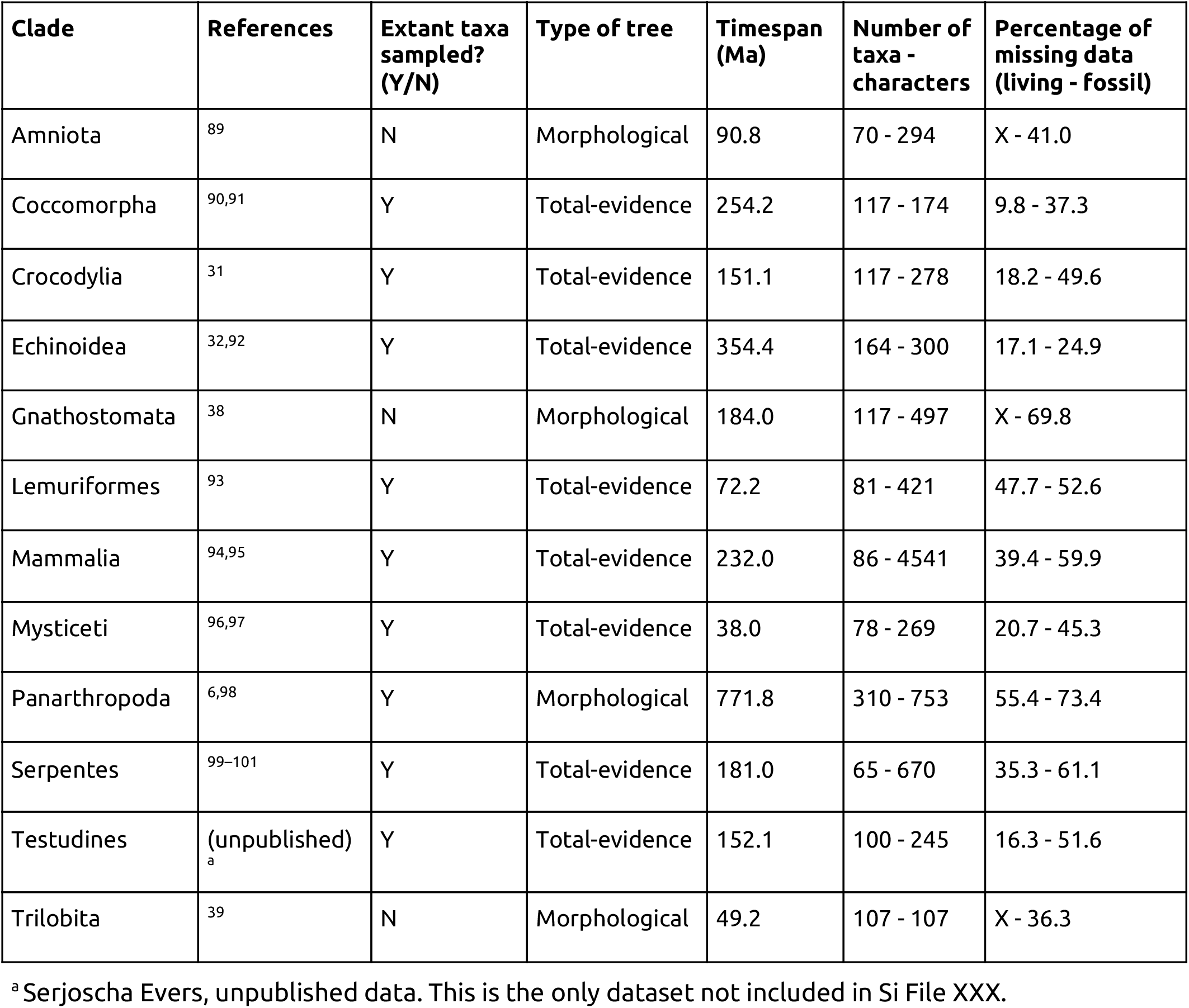
Empirical datasets used to determine realistic properties of our simulated datasets (see Figs. S5-6). For each clade, citations correspond to studies who contributed morphological matrices, tree topologies, or both. Some phylogenies employed only morphological data, others combined morphology and molecular evidence under total-evidence approaches, all were inferred using tip-dated methods.

## Supplementary Figures

**Figure S1:**
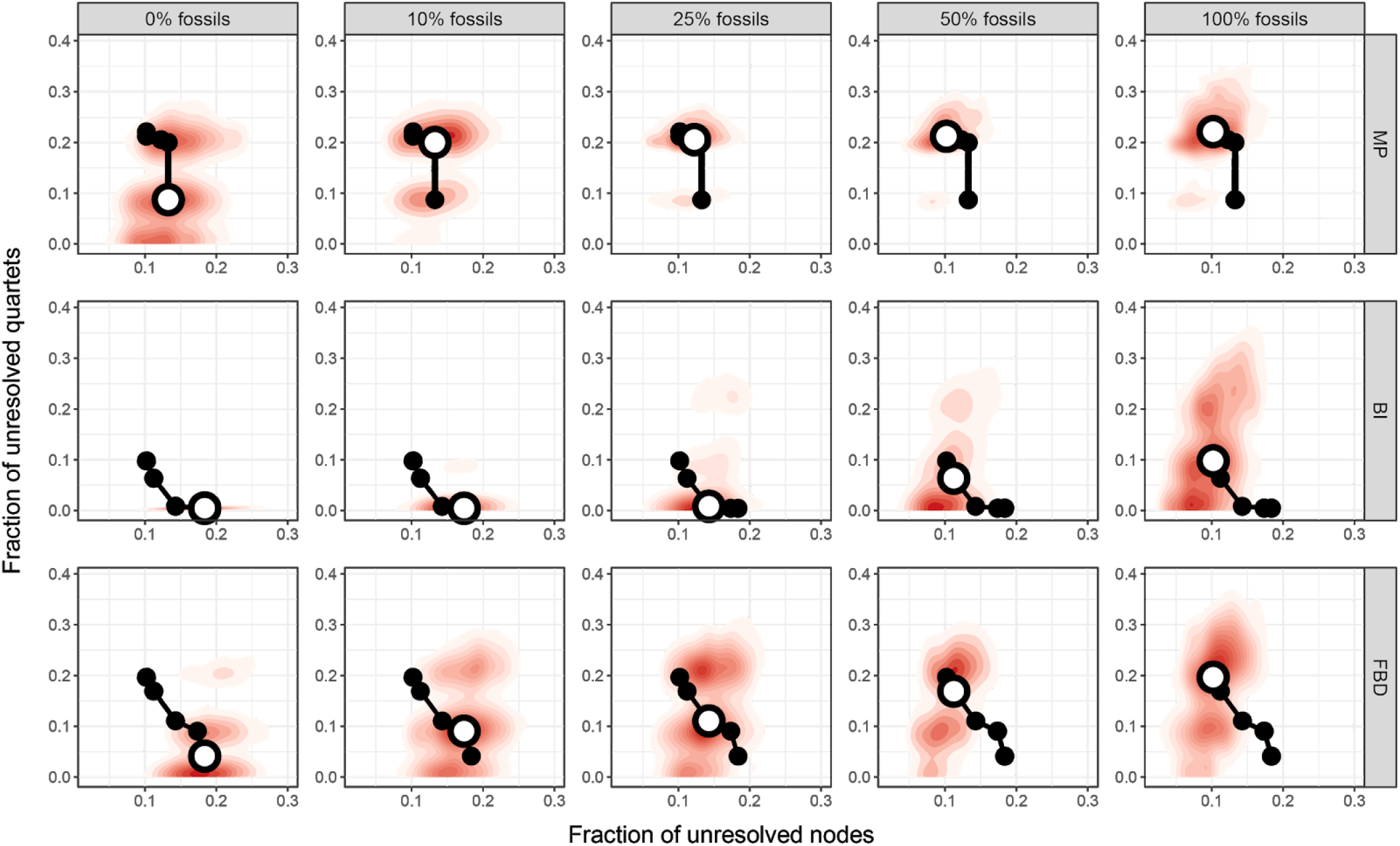
Relationship between quartet and bipartition-based precision. Red contour lines show the kernel density of all consensus topologies across different conditions. Circles show the median of the distributions as fossil sampling increases, with the median for the corresponding quadrant highlighted in white. Only data for the condition of no missing data are shown. The increased sampling of fossil terminals decreases the fraction of unresolved nodes, while at the same time increasing the fraction of unresolved quartets; i.e., topologies are more resolved, yet are more likely to exhibit unresolved deep nodes (see also Fig. 3).

**Figure S2:**
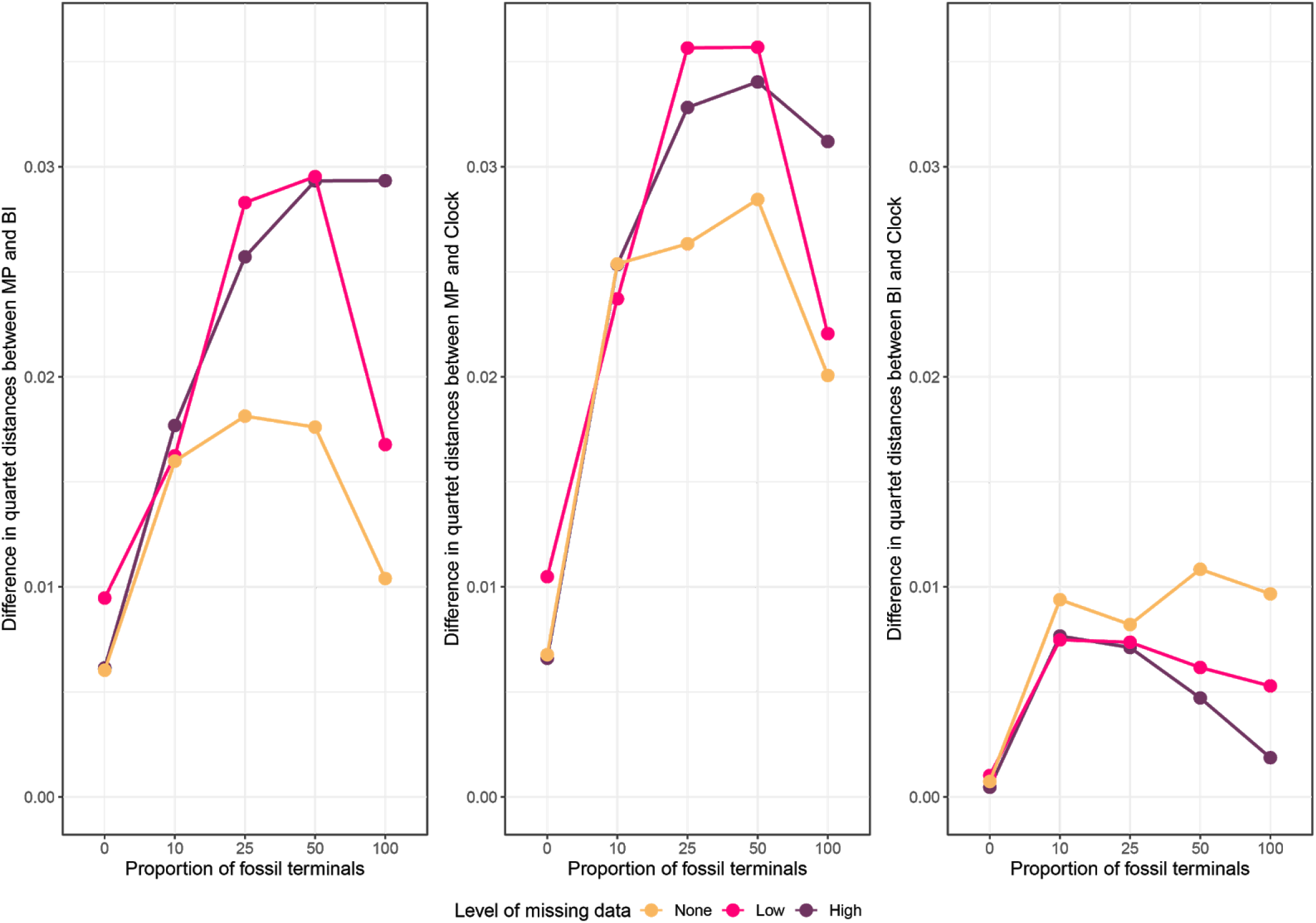
Difference in performance (average quartet distance) between pairs of inference methods across conditions of missing data. Quartet distances of the better-performing method are subtracted to that of the worse-performing method, rendering positive absolute differences. The difference in performance between MP and probabilistic methods widens with increasing missing data, showing part of the success of probabilistic methods is their robustness to missing data. On the other hand, the benefits of tip-dating decrease with missing data.

**Figure S3:**
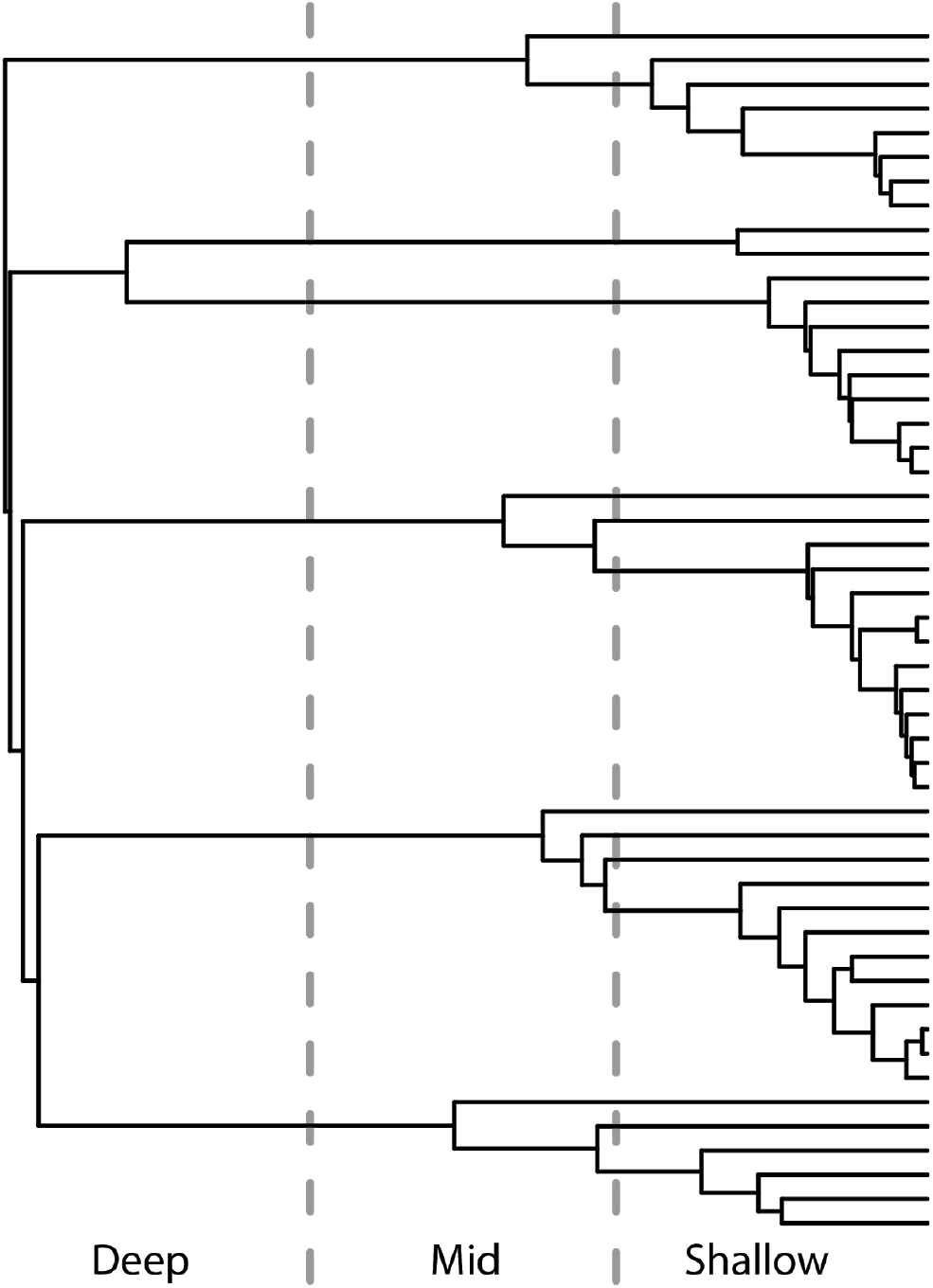
Example of how nodes connecting extant taxa were binned into three time-slices of equal duration. Topology corresponds to a randomly selected simulated tree, pruned to retain only the 50 extant terminals present in a replicate that sampled 50% fossil tips. The ‘deep’ category includes the nodes involved in the simulated event of radiation, the mid category incorporated most of the earliest divergences within each of the main clades, while the shallow category includes most divergences within these clades.

**Figure S4:**
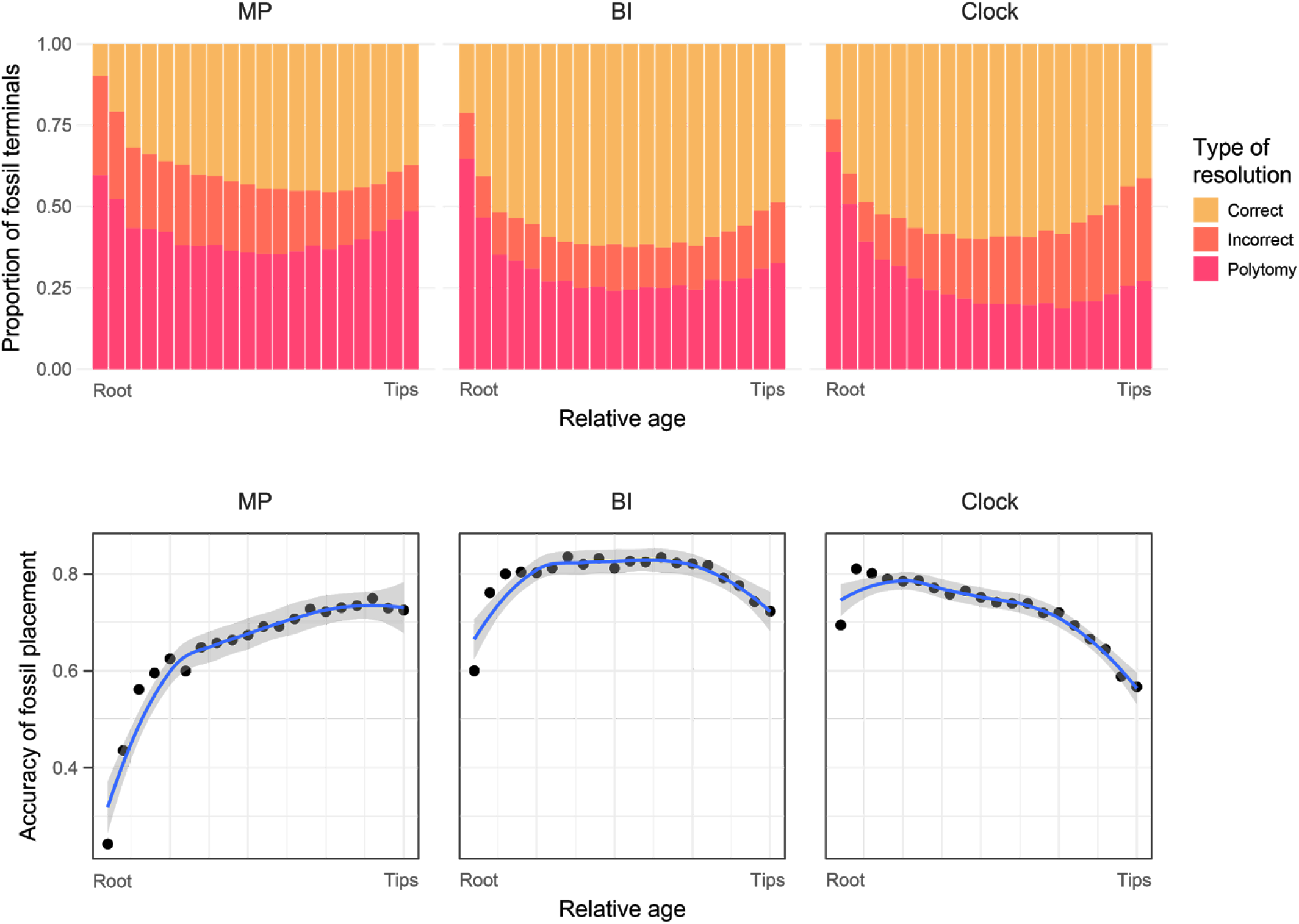
Effect of age on the resolution of fossils across inference methods. Results correspond to all fossils across datasets with high levels of missing data. Fossil terminals are binned into 20 time-slices of equal duration, spanning the time from root to tips. **Top:** Type of resolution (correct, incorrect, unesolved) of fossil terminals as a function of age. MP exhibits a comparatively low proportion of correct placements compared to probabilistic methods. Tip-dated Bayesian inference has the highest proportion of incorrectly resolved young fossils of all methods. **Bottom:** Accuracy of fossil placement as a function of age, measured as the ratio between the proportion of correctly resolved fossils divided by the proportion of resolved fossils. Tip-dated inference exhibits a low accuracy in the placement of younger fossil terminals.

**Figure S5:**
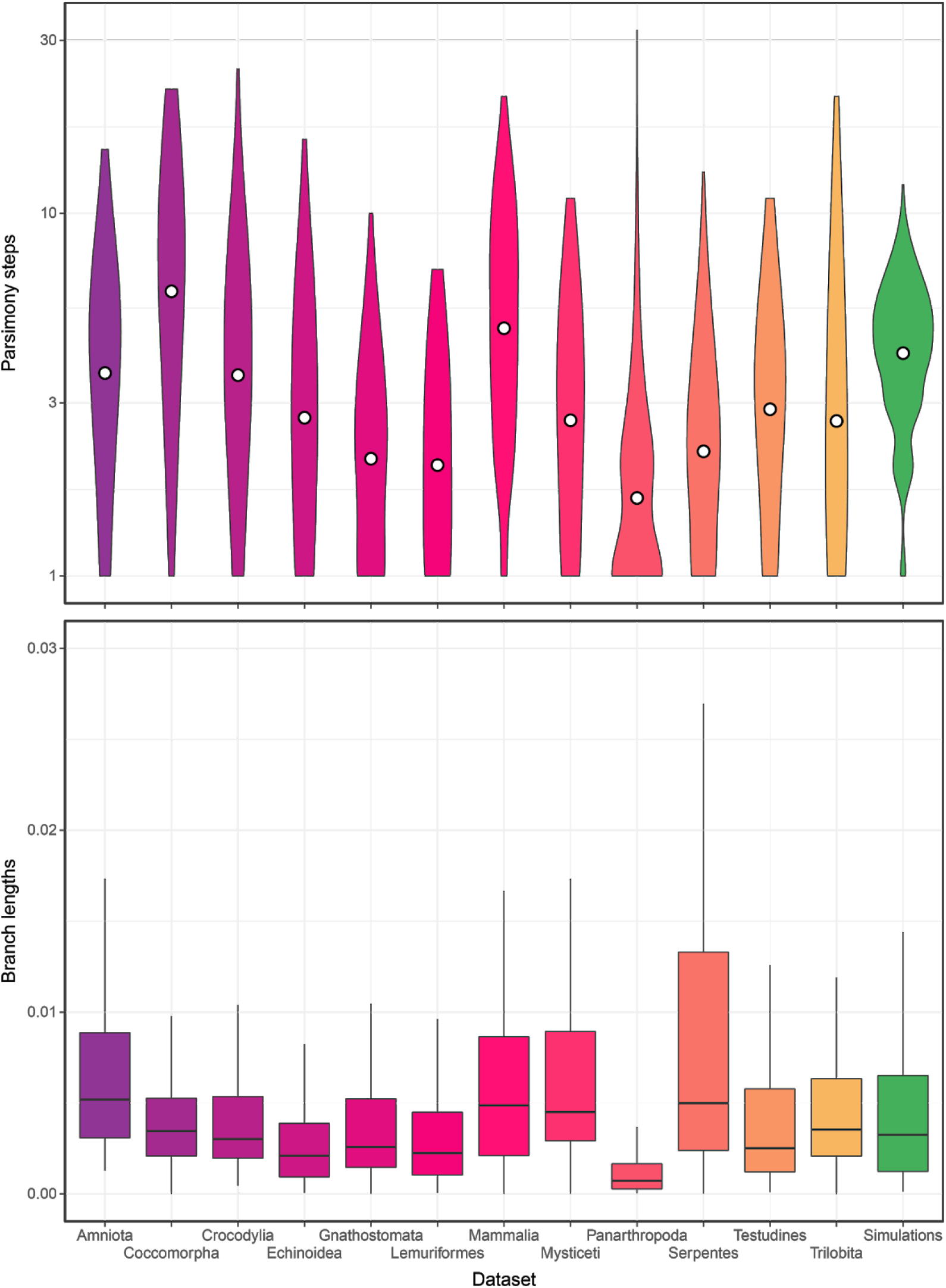
Comparison of 25 randomly-sampled simulated datasets against empirical data. High (realistic) levels of missing data were input, and simulated datasets were pruned to 100 randomly-selected taxa and 300 parsimony-informative characters, as this is the size of datasets analyzed. **Top**. Number of parsimony steps, as a proxy for overall morphological rate and variability. Empirical datasets were reduced to only binary, parsimony-informative characters, to provide a better comparison to the type of characters generated by TREvoSim. **Bottom**. Distribution of branch lengths, expressed as proportion of total tree length. For both metrics, simulated values are within 95% confidence intervals made from the combined empirical distributions, and mean values of simulated datasets are contained within the range of mean values for empirical datasets.

**Figure S6:**
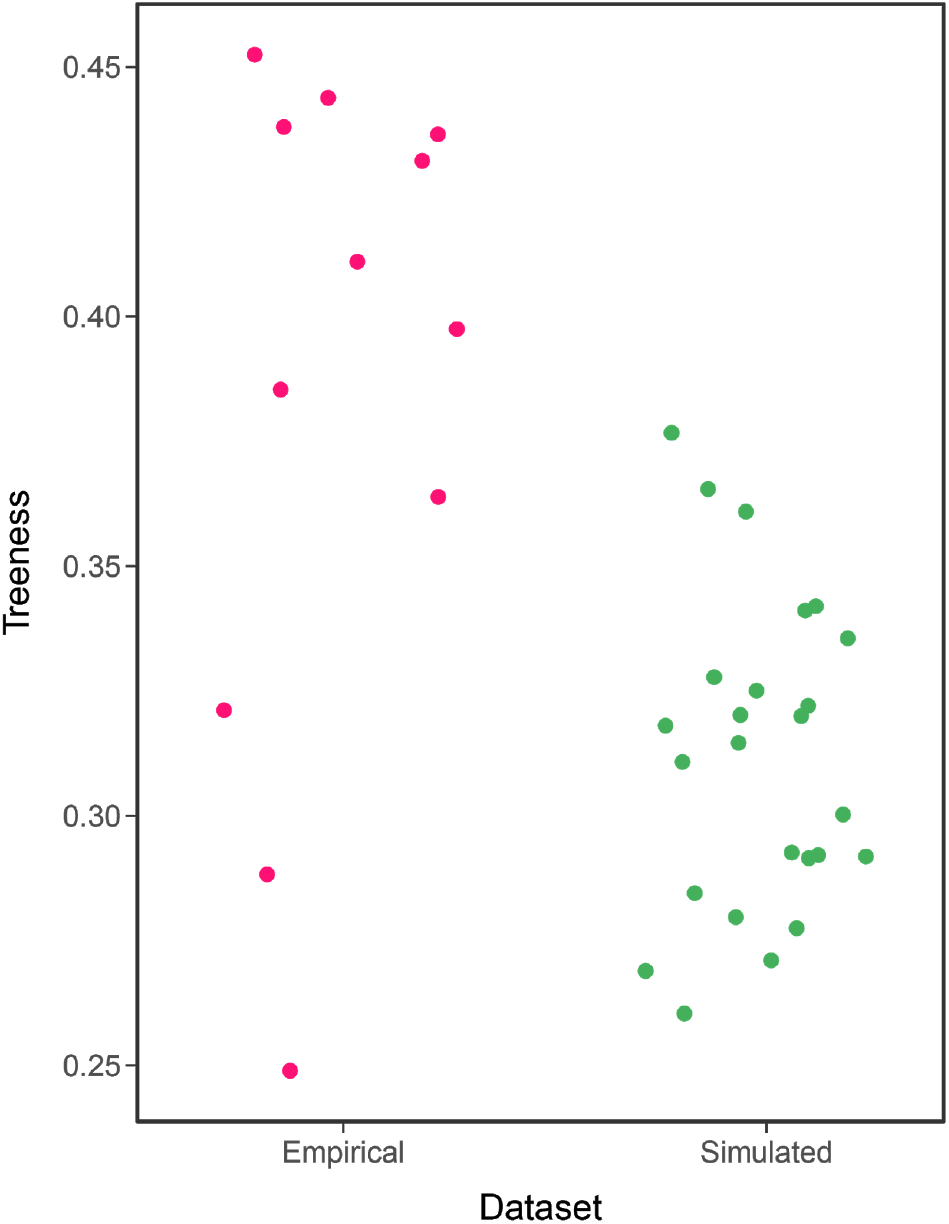
Comparison of 25 randomly-sampled simulated datasets against empirical data with respect to their levels of treeness. Simulated datasets were pruned to 100 randomly-selected taxa. Treeness represents the fraction of total tree length that is on internal branches^44^. Simulated datasets fall within the range of empirical datasets.

**Figure S7:**
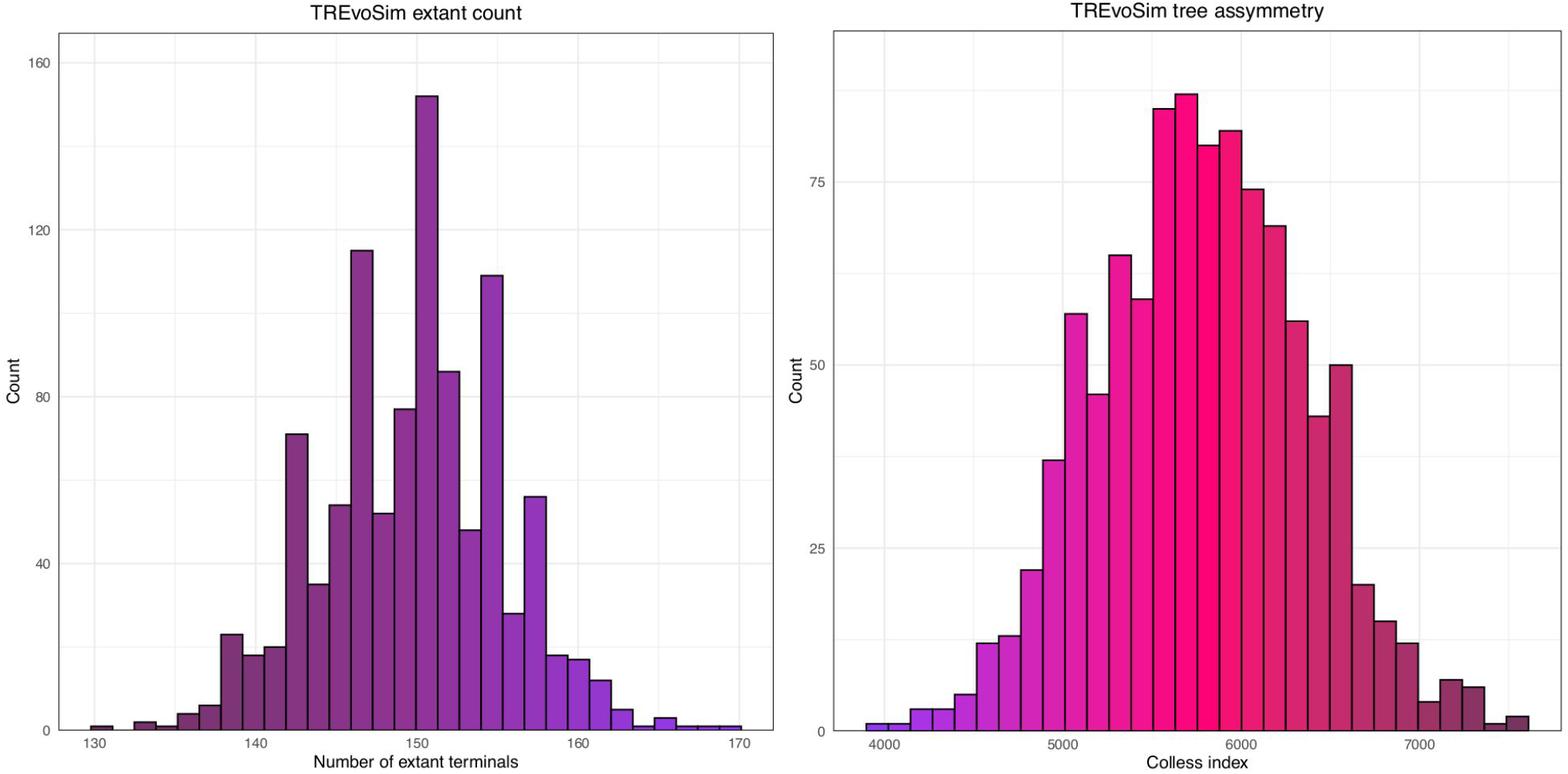
Simulated topologies exhibit a range of values of extant terminals (left) and tree symmetry (right), measured using Colless’ index^56^. Mean number of extant terminals (out of a total of 999) = 149.6, range = 136-169.

**Figure S8:**
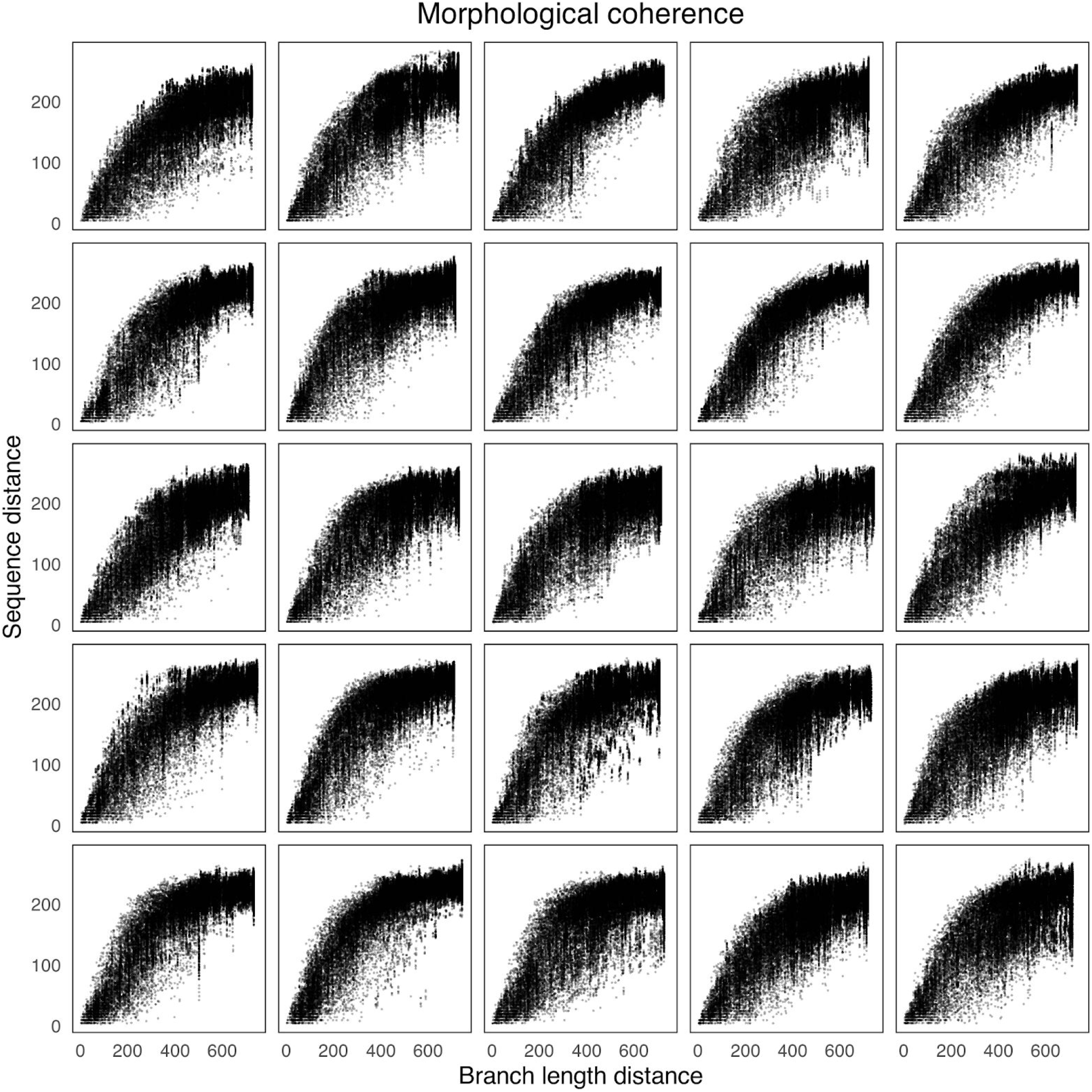
Morphological coherence^17,58^ of 25 randomly chosen simulations. Morphological and patristic distances show high levels of correlation, with signs of morphological saturation only at among the most distantly-related terminals.

**Figure S9:**
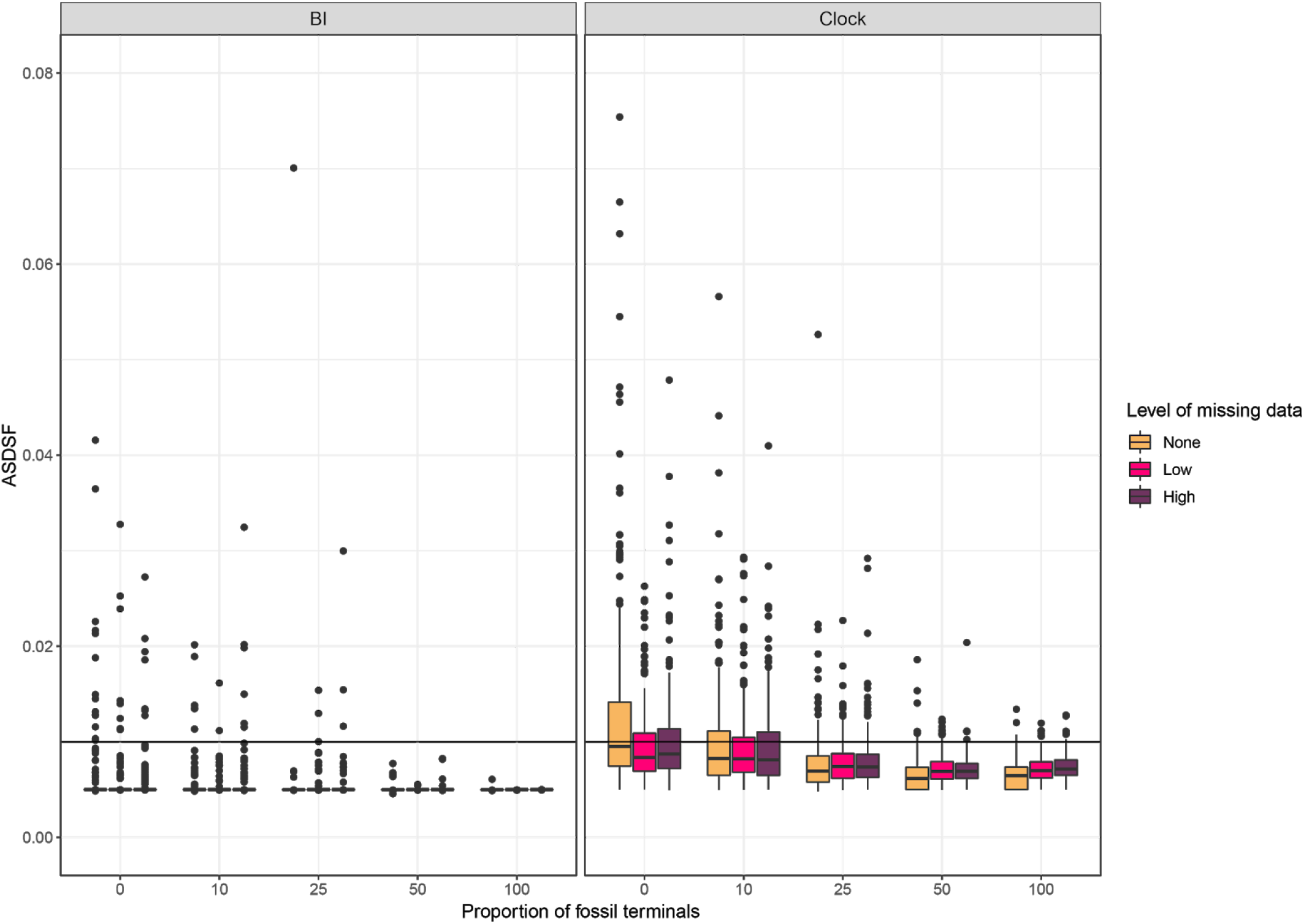
Topological convergence of Bayesian runs (undated and tip-dated). The average standard deviation of split frequencies (ASDSF) of runs was registered upon completion to check that the requested number of generations was sufficient to adequately sample from the posterior distribution. The median ASDSF value for each condition of missing data and fossil sampling is below 0.01 (horizontal line) across BI and Clock analyses. This threshold was used by previous studies^9,18^ to terminate runs, considering it represented an accurate sampling of the posterior distribution of topologies. The majority of analyses performed here were continued until ASDSF values were even lower. Increased fossils sampling helps attain convergence under both tip-dated and undated Bayesian inference.

